# Cooperation Destabilizes Communities, but Competition Pays the Price

**DOI:** 10.1101/2023.12.26.573354

**Authors:** Ananda Shikhara Bhat, Suryadeepto Nag, Sutirth Dey

**Affiliations:** Department of Biology, Indian Institute of Science Education and Research, Pune, Maharashtra - 411008, India; Institute of Organismic and Molecular Evolution, Johannes Gutenberg University Mainz, Hanns-Dieter-Hüsch-Weg 15, 55128 Mainz, Germany; Faculty of Geosciences and Environment, Institute of Geography and Sustainability, University of Lausanne, Geopolis, Mouline, CH-1015 Lausanne, Switzerland

**Keywords:** Cooperation, Individual-based model, Interspecific interactions, Ecological stability, Community ecology, Lotka-Volterra, Coexistence

## Abstract

A classic result in theoretical ecology states that an increase in the proportion of cooperative interactions in unstructured ecological communities leads to a loss of stability to external perturbations. However, the fate and composition of the species that constitute an unstable ecological community following such perturbations remains relatively unexplored. In this paper, we use an individual-based model to study the population dynamics of unstructured communities following external perturbations to species abundances. We find that while increasing the number of cooperative interactions does indeed increase the probability that a community will experience an extinction following a perturbation, the entire community is rarely wiped out following a perturbation. Instead, only a subset of the ecological community is driven to extinction, and the species that go extinct are more likely to be those engaged in a greater number of competitive interactions. Thus, the resultant community formed after a perturbation has a higher proportion of cooperative interactions than the original community. We show that this result can be explained by studying the dynamics of the species engaged in the highest number of competitive interactions: After an external perturbation, those species that compete with such a ‘top competitor’ are more likely to go extinct than expected by chance alone, whereas those that are engaged in cooperative interactions with such a species are less likely to go extinct than expected by chance alone. Our results provide a potential explanation for the ubiquity of cooperative interactions in nature despite the known negative effects of cooperation on community stability.

## Introduction

One of the central questions in community ecology asks whether observed patterns of interspecific interactions can be explained using ecological principles (Dobson et al., 2020; Sherratt et al., 2009; Sutherland et al., 2013). An important notion in this regard is the ‘stability’ of a community i.e., its ability to withstand external disturbances, and how it is affected by interactions between the constituent species (Sherratt et al., 2009). While the effects of interaction patterns on community stability have been extensively studied in the literature (Allesina and Tang, 2012; Coyte et al., 2015; May, 1973; Qian and Akçay, 2020; Serván et al., 2018; Stone, 2020; Mougi and Kondoh, 2012; Mougi, 2016a; Mougi, 2016b; Mougi and Kondoh, 2016), the question of what happens to unstable communities following a perturbation has received much lesser attention. Furthermore, the ecological literature has used a plethora of stability concepts (Grimm and Wissel 1997). While stability in deterministic systems can reasonably be construed as the ability of a population to return to its exact previous configuration following a perturbation (‘resilience’ s*ensu* Grimm and Wissel 1997), this may be too restrictive of a criterion for finite, stochastic communities. A more natural stability criterion for stochastic systems is ‘persistence’, i.e. the ability of a community to retain all its species following a perturbation (Grimm and Wissel 1997, Jansen and Sigmund 1998). In this paper, we ask how stability in the sense of persistence is affected by the amount of mutualism present in a community. Furthermore, if a community is unstable in the persistence sense, we ask whether all the species in a community go extinct after a perturbation, and if not, whether the composition of the community in terms of the interaction patterns is changed following species loss.

The interactions between two species in a community can be broadly classified into three types: competitive, cooperative, and exploitative (Allesina and Tang, 2012). When two competing species interact, both are adversely affected and perform worse than if the other were absent. This could be due to several reasons, including competition for shared resources or the secretion of toxins that harm members of the other species. Contrastingly, cooperative species help each other grow, for example, by providing each other with essential resources. Finally, exploitation refers to the phenomenon where one of the two interacting species benefits from the interaction, whereas the other is harmed. Such interaction patterns would biologically correspond to phenomena such as predation or parasitism, in which individuals of one species actively consume part or whole of individuals of the other species (Allesina and Tang, 2012).

Modeling all species interactions in a community through pairwise interactions (as in Lotka-Volterra type models), interspecific effects can be collected in a so-called ‘interaction matrix.’ The *i-j*^th^ entry of the interaction matrix quantifies the effect of species *j* on the growth rate of species *i*. This, of course, need not be exclusive to pairwise interactions alone, as higher-order interactions among multiple (greater than 2) species may be decomposed into multiple pairwise interactions. Diagonal entries of such a matrix capture the strength of intraspecific competition. If we assume, as is often done (e.g. (Allesina and Tang, 2012; Coyte et al., 2015; May, 1973)), that interactions within the community are completely random (unstructured), this matrix can be entirely characterized by measuring the fraction of cooperative (*p*_*m*_), competitive (*p*_*c*_), and exploitative (*p*_*e*_) interactions present. Under such a scenario, the question of how community interaction patterns affect stability reduces to how *p*_*m*_, *p*_*c*_, and *p*_*e*_ affect stability.

Previous theoretical studies in the infinite species richness limit have shown that if interactions are mostly pairwise and linear (i.e. modelled by Lotka-Volterra like dynamics) then for a given magnitude of intraspecific competition, unstructured communities with a greater fraction of cooperative interactions (higher pm) are less likely to be stable (Allesina and Tang, 2012; Coyte et al., 2015). Here, the concept of stability used is ‘resilience’ (Grimm and Wissel 1997), i.e. whether a system can return to the same initial configuration of species abundances following an external perturbation. These results were extended to a system with finite (but large) species richness using a computational IBM (Coyte et al., 2015). The latter study also used a different stability concept called ‘persistence’ (Grimm and Wissel 1997), i.e. the ability of a community to retain all its species following a perturbation. Coyte et al. 2015 showed that communities with higher *p*_*m*_ are more likely to lose species to extinction following random perturbations to their abundances, and thus that mutualism is also destabilizing when persistence is used as a stability concept. Since environmental conditions are seldom constant, perturbations are ubiquitous in nature. If communities with a large fraction of cooperative interactions are unstable, and unstable communities are more likely to experience species loss, cooperative communities would be less likely to be found in nature (but see (Bascompte and Scheffer, 2023; Bastolla et al., 2009; Holland et al., 2002; Lever et al., 2020; Ringel et al., 1996) for how these expectations are altered by community structure or density dependence). However, this insight is difficult to reconcile with the empirical observation that many communities with reasonable number of positive interactions (cooperation and commensalism) are abundant in ecological communities (Kehe et al., 2021), with some communities almost solely comprising of cooperative interactions (Machado et al., 2021).

Another issue in terms of community composition relates to the relative strengths of the intra-specific and inter-specific interactions. Theoretical studies (May, 1973; Allesina and Tang, 2012; Barabás et al., 2016, 2017) suggest that species-rich, randomly assembled communities are overwhelmingly likely to be unstable (in the resilience sense) unless intraspecific competition is much stronger than interspecific competition. This insight is congruent with the observation that the interaction networks of large microbial communities in nature often have low connectivity (Yonatan et al., 2022). If many species-rich communities are likely to be unstable, several questions arise related to their fate. For example, if an external perturbation causes some extinctions in a randomly assembled community, do all species go extinct? If not, then is the identity of the species that go extinct random, or determined by their interactions with other members of the community?

In this paper, we study the effects of interspecific interactions on both community persistence as well as resultant community composition in the face of external perturbations. We employ a stochastic individual-based model (IBM) in which we supply rules at the species level and let population-level patterns emerge spontaneously. Our IBM follows Lotka-Volterra growth patterns except for imposing an additional cap on species numbers (due to limited resources). This cap allows us to relax the unrealistic assumption of infinite unbounded growth rates associated with cooperative interactions in purely analytic Lotka-Volterra models (Allesina and Tang, 2012) to study the effects of cooperation in more realistic communities. While we focus on unstructured communities for the rest of this article, our approach is general and can easily be extended to analyze communities with arbitrary interaction structures. Since our system is inherently stochastic, we study stability in the persistence sense, asking whether the probability of losing a species to extinction following a perturbation depends on community structure (Coyte et al., 2015), and if so, whether the species lost are random with respect to the interactions they are engaged in. Our primary results indicate that perturbations to unstable communities do not necessarily result in all species of the community going extinct. Instead, we find that perturbations often result in the extinction of only a few species and the formation of a new “sub- community” with fewer species. We observe that while communities with a greater fraction of cooperative interactions are more likely to experience extinctions of some species, the resultant community formed after the extinctions is found to have a greater fraction of cooperative interactions than the original community, implying that species which partake in competitive interactions are more likely to be selectively removed. We then study the community dynamics to answer why species participating in competitive interactions are more likely to go extinct.

## Methods

### Model Overview

We present a brief overview of the model here, with all the relevant details in the following subsections.

This study uses an Individual-Based Model based on a previous framework (Coyte et al., 2015). In this framework, every species interacts with every other species either cooperatively, competitively, or exploitatively. We begin with a community consisting of *S* species. For the rest of the manuscript, we use either *S=7* or *S=15*. Let *p*_*m*_, *p*_*c*_, and *p*_*e*_ denote the fraction of total cooperative, competitive, and exploitative interactions in the community respectively. Following usual conventions (Allesina and Tang, 2012; Coyte et al., 2015; May, 1973; Serván et al., 2018), we assume that all species have the same level of intraspecific competition, denoted by -*s*. To study the effects of species interactions on persistence, we initialize a community with a specified (*p*_*m*_, *p*_*c*_, *p*_*e*_) at equilibrium determined from Lotka-Volterra dynamics. To do this, we generate an interaction matrix such that a randomly drawn interspecific interaction will be cooperation with probability *p*_*m*_, competition with probability *p*_*c*_, and exploitation with probability *p*_*e*_. Interaction strengths are drawn from a gamma distribution. We chose a Gamma distribution arbitrarily, since previous work has shown that only the first two moments of the distribution affect the results, while the precise choice of distribution (Gamma, half-Normal, etc.) is irrelevant (Coyte et al., 2015). We then initialize the community with a random initial configuration of species abundances, and adjust the growth rates of each species such that this initial configuration is a fixed point of the generalized Lotka-Volterra equations. We then perturb this community by killing off a fixed, relatively small number of randomly chosen individuals (for details, see below). The strength of this perturbation was sufficiently small such that no species went extinct due to the perturbation alone. Thus, any subsequent species extinctions would be driven by the dynamics of biotic interactions within the community following the perturbation. We observe how species abundances vary following such a perturbation by allowing the community to stabilize to a new steady state according to the generalized Lotka-Volterra equations.

Figure 1 provides a schematic description of our model. To avoid the possibility of only observing the effects of specific network topologies, we randomize the specific interactions between species within a community between replicates such that only the effects of the proportion of each interaction type (as opposed to the specific network topology) were kept constant. Since an extinction in a community reduces the species richness of the community and thus reduces the number of interactions, we define a new interaction matrix following extinctions, where the interaction effects from the extinct species have been set to 0. The effects of any potential extinctions on community composition can then be studied by comparing the species richness and the interaction matrix at the beginning and the end of the simulation. For simplicity, we present results for *p*_*e*_ = 0 (i.e., communities where all interactions are cooperative or competitive) in the main text. Results for communities containing exploitative interactions (*p*_*e*_ > 0) are presented in the supplementary information (Figs S1-S7).

**Figure 1:**
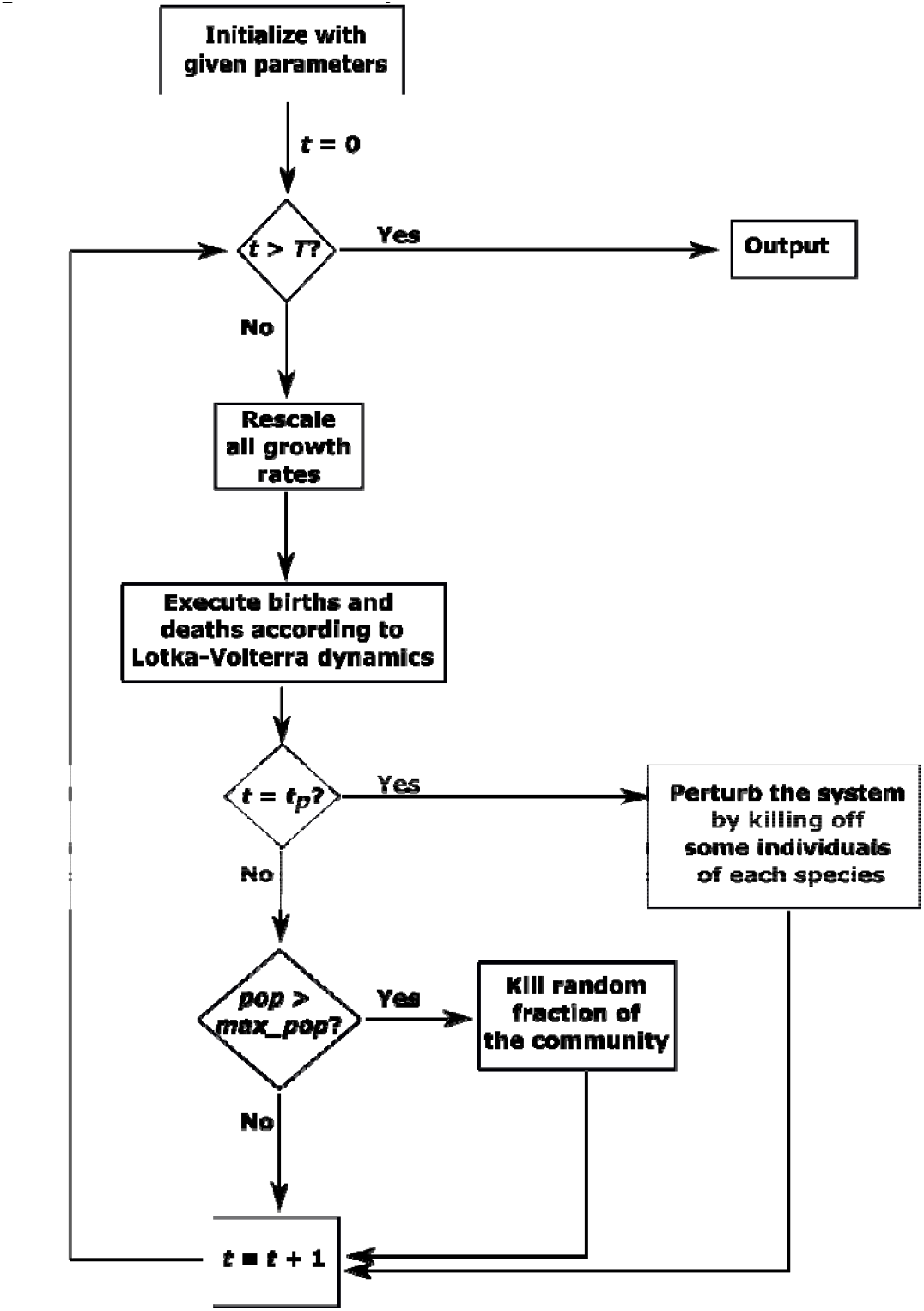
A schematic description of our IBM. The model is initialized with a set of parameters and a time counter set at t=0. In this diagram, t is the number of time steps in the model (Not to be confused with the actual time for which the dynamics are simulated; the two differ due to adaptive rescaling of the growth rates, see Methods section). The model is allowed to run for a total of T timesteps. The community follows Lotka-Volterra dynamics, except when t_p_ timesteps have elapsed, at which point the community experiences an external perturbation that kills off some individuals of each species. If at any time the Lotka-Volterra dynamics predict that the total population size of the community exceeds a maximum allowed size, a random fraction of the community is eliminated until the total size is below the maximum allowed size. Refer to the main text for details of the model and the parameter values chosen.

### Initialization and static parameters

The world of the simulation is a 50×50 2D square lattice with periodic boundary conditions. Each lattice site is occupied by, at most, a single individual. Every individual interacts with every other individual, and thus interactions are non-local. An individual can only reproduce if an empty space is adjacent to the focal individual. The community is initialized with *S* different species, all interacting with each other (this corresponds to setting the connectivity *C = 1* in the Coyte *et al*. 2015 IBM). The total population size is capped at 90% lattice occupancy at any given time. Thus, our simulations could have a maximum of 2250 individuals coexisting at a given time in a given simulation run. An obvious drawback of this design is a limit on how many species we can include in our community. However, highly species-rich communities require each species to have high intraspecific competition, at times orders of magnitude greater than the other interactions, to ensure stability of the community as a whole in either the resilience or the persistence sense (May, 1973; Allesina and Tang, 2012; Barabás et al., 2016, 2017; Bascompte and Scheffer, 2023; Coyte et al., 2015; Serván et al., 2018). Species-rich communities in which interspecific interactions are of the same order as intraspecific interactions are therefore inherently unstable, and studying the effects of the distribution of interaction patterns (cooperation, competition, and exploitation) in such communities is thus of limited biological use.

### Interactions and species abundance dynamics

Species abundance dynamics are assumed to follow the generalized Lotka-Volterra equation, which, in matrix form, reads

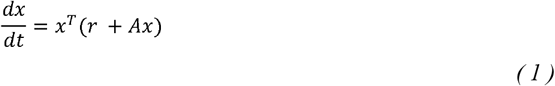

For a community with *S* species, equation (1) is an *S-*dimensional equation. Here, *r* is an *S*-dimensional vector of intrinsic growth rates, and the interaction matrix *A* is an *S × S* matrix that captures the effects of interspecific interactions. For our simulations, we use either *S = 7* or *S = 15*. The *ij*-th entry of the matrix, which we denote by *a*_*ij*_, describes the effect of species *j* on the growth rate of species *i*. Diagonal entries of this matrix are *-s*, which is a parameter that controls the strength of intraspecific competition. For off-diagonal entries, the magnitude of *a*_*ij*_ is drawn from a Gamma distribution with a mean of 0.25 and a variance of 0.01. Since the probability of drawing a value that is exactly 0 from such a continuous distribution is extremely small, the connectance of our communities is very close to 1 (i.e. every species interacts with every other species) across all simulations. Since we wish to model an unstructured community, the signs of *a*_*ij*_ are determined randomly such that species *i* and *j* have a cooperative interaction (+/+) with probability *p*_*m*_, have a competitive interaction (-/-) with probability *p*_*c*_, and an exploitative interaction (+/-) with probability (1- *p*_*m*_ - *p*_*c*_). In the case of an exploitative interaction, each species is equally likely to be the one that is benefited. For all simulations in the main text, we set *p*_*m*_ and *p*_*c*_ such that *p*_*m*_*+p*_*c*_*=1*. Thus, there were no exploitative interactions in the community for results presented in the main text. Since we fill the interaction matrix according to probabilistic rules, *p*_*m*,_ *p*_*c*,_ and *p*_*e*_ are not the realized values in any given simulation but are the parameters used to determine the probability of an interaction being cooperative, competitive, or exploitative. Note that since the sign of each entry of the matrix is independently drawn, a single species may be engaged in multiple types of interactions. When initializing the system, each species is assigned a random population density, and the growth vector *r* is chosen such that the population densities at initialization are a fixed point for the Lotka -Volterra dynamics. Thus, given a realized interaction matrix *A* and a randomly initialized population vector *x*_*0*_, we set the growth rate to

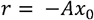

This ensures that the RHS of Eq. (1) becomes zero at the initial community configuration *x*_*0*_ and the community thus begins at equilibrium.

Since species growth dynamics are modelled entirely by Eq. (1), spatial structure does not affect the interaction dynamics in our model. Instead, the lattice serves to impose a population cap and prevent unbounded growth.

This constraint is meant to reflect the fact that in ecological communities, resources can be limited, preventing unbounded growth, without interaction potential being limited.

### Perturbation and extinctions

Following a previous study (Coyte et al., 2015), we implement a perturbation five time steps into the simulation by eliminating 10% of the population of each species. The individuals eliminated are chosen at random. We chose a value of 10% because we wished to investigate the impact of perturbations that affect the population dynamics (biologically corresponding to relatively minor environmental fluctuations) but are not so large as to introduce bottleneck effects or cause a species to go extinct due to the perturbation alone. If larger perturbation strengths are used, stochastic effects may dominate the dynamics. The simulation is then allowed to run until *T*=750 time steps have passed. For consecutive perturbations, as depicted in Figure 1C, we execute a secondary perturbation at 750 time steps and allow the system to run for an additional 750 time steps (*i*.*e*., for a total time of *T*=1500). These particular time steps are chosen as the population attained an equilibrium well before this time in sample simulations. If, at any point in the simulation, the growth rates predicted a population size that exceeded the maximum population size allowed by the simulation, a random fraction of the population is killed off such that the new population size is below the maximum allowed population size. Thus, unlike in classic analytical Lotka-Volterra models of cooperation (Allesina and Tang, 2012; Coyte et al., 2015), species in our IBM cannot exhibit infinite unbounded growth due to limited space.

### Computing realized growth rates following a perturbation and an adaptive timestep

Following the perturbation, the community is no longer at the density *x*_*0*_ and is thus no longer at equilibrium for the Lotka-Volterra dynamics defined by Eq. (1). While computing the resultant population dynamics, we rescale the growth rates to allow for an adaptive timestep using a method introduced in a previous study (Coyte et al., 2015). We first define a parameter *g*_*cap*_ which is the maximum magnitude of growth rate allowed in a single time step. We then rescale the realized growth rate of every species to be in [-*g*_*cap*_, *g*_*cap*_]. This rescaling lets us naturally define an adaptive time step that enables the model to simulate long times if differences in growth rates between species are large, while also allowing us to determine the fine-scale dynamics if the differences in growth rates is small (Coyte et al. 2015). More explicitly, our model can be described as follows: At time τ, we first compute the realized per-capita growth rate of species *i, g*_*i*_, as

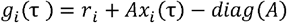

The *-diag(A)* term simply serves to remove the effect of an individual on itself. We then find the maximum growth rate (in absolute value), *g*_*max*_(*τ*) = *max* |_*i*_ *g*_*i*_(*τ*)|, and rescale the growth rate of each species as

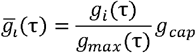

We then advance time according to the formula

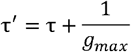

The population density of species *i* at time *τ*^*′*^ is then given by

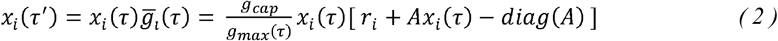

This constitutes one ‘time step’ of the model. Thus, though the model runs for only 750 (or 1500) time steps, the true ‘in world’ time is considerably greater than 750 (or 1500), since time is rescaled dynamically at every step. The extent of the difference between time steps of the model and simulated time within the model is controlled both by the parameter *g*_*cap*_ and by the difference between the realized growth rates *g*_*i*_ of the different species. Since the right-hand side of equation (2) is just a rescaled version of the right-hand side of equation (1), introducing adaptive time steps in this manner only rescales time and does not otherwise affect the behavior of the system (Coyte et al. 2015). Such adaptive time-stepping greatly reduces the computational resources required to simulate the model.

### Computing interaction structures following extinctions

We do not modify the interaction matrix *A* in any way following species interactions since extinctions already affect the growth dynamics in Eq. (1) through the density vector *x*. At the end of the simulation, we check which species are extant (*i*.*e*. have density *x*_j_ > 0) and compute the fraction of cooperative interactions (*p*_*m*_) and competitive interactions (*p*_*c*_) in the new community formed solely of extant species. These new fractions let us examine whether extinctions are non-random with respect to initial interactions (*i*.*e*. whether species engaged in more cooperative interactions are more or less likely to go extinct than those engaged in more competitive interactions).

### Software and statistical analysis

Since our simulations are stochastic, we run 100 independent realizations of the simulation for any given combination of parameter values to obtain an estimate of expected (average) behavior. We vary *p*_*m*_ from 0 to 1 (this automatically varies *p*_*c*_ from 1 to 0) and *s* from 0.15 to 1.65. All parameters that are held constant for all simulations run in this paper are summarized in Table 1. All simulations were run in Python 3.6. Plots were made using either Python 3.11.4 (packages numpy 1.24.2, pandas 1.5.3, matplotlib 3.7.1, and seaborn 0.12.2) or R 4.2.2 (packages dplyr 1.1.1, rstatix 0.7.2, ggplot2 3.4.1, and ggpubgr 0.6.0). Statistical tests were run in R 4.2.2. For effect size calculations, we used the *wilcox_effsize* function from the rstatix package.

**Table 1:**
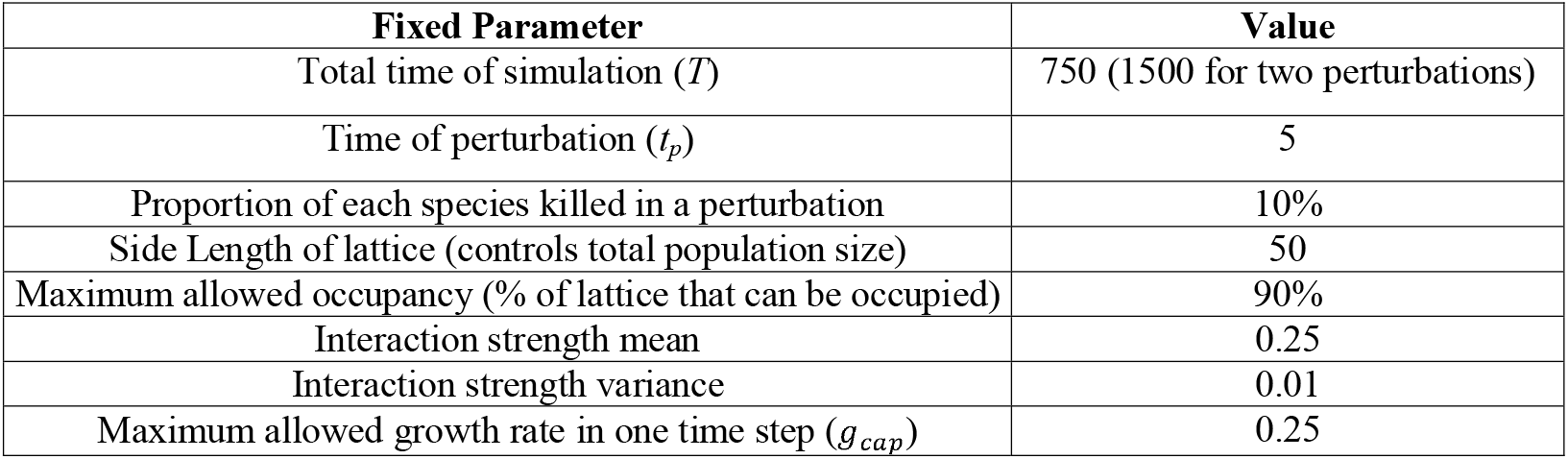
Parameters that were held constant across all simulation runs. The interaction strengths were drawn from a Gamma distribution with the specified mean and variance.

| Fixed Parameter | Value |
| --- | --- |
| Total time of simulation ($T$) | 750 (1500 for two perturbations) |
| Time of perturbation ($t_p$) | 5 |
| Proportion of each species killed in a perturbation | 10% |
| Side Length of lattice (controls total population size) | 50 |
| Maximum allowed occupancy (% of lattice that can be occupied) | 90% |
| Interaction strength mean | 0.25 |
| Interaction strength variance | 0.01 |
| Maximum allowed growth rate in one time step ($g_{cap}$) | 0.25 |

## Results

To briefly recapitulate our modeling approach: we use an IBM to study the stability and dynamics of finite- species communities after perturbations. We consider communities with 7 or 15 species interacting with every other species through cooperation or competition. Our model also includes intra-specific competition, i.e., when members of each species compete with conspecifics. Further, since the total amount of space is limited, our IBM cannot exhibit infinite unbounded growth, a well-known unrealistic prediction of analytical Lotka-Volterra models of cooperation (Allesina and Tang, 2012; Coyte et al., 2015). We initiate communities in equilibrium and simulate a perturbation, where a fraction of all community individuals is killed off randomly. We then study the stability and composition of the post-perturbation community. If the original community were stable, the species composition would remain unchanged after perturbation.

### Unstable communities do not usually lose all their species following a perturbation

In our study, we perturb the communities in such a way that the strength of the perturbation in itself does not lead to species extinctions. Under such a scenario, the community does not lose its entire assemblage of species to extinctions but instead forms a smaller community with fewer species (Fig 2A). A recent study has also independently reported the same result using numerical simulations of Lotka-Volterra dynamics (Rohr et al., 2025). We also found that following a perturbation, more species-rich communities (Fig 2A. compare red and yellow curves) and communities with a greater fraction of cooperative interactions (i.e., higher pm) (Fig 2A, 2B) tended to lose more species, recapitulating classic results based on resilience stability (May, 1973; Allesina and Tang, 2012). Thus, we can say that mutualism is destabilizing in our work, since a community loses more species as *p*_*m*_ increases. However, the entire community went extinct only when the fraction of cooperative species was very high (Fig 2B). In line with previous studies based on resilience, we found that increasing intraspecific competition promoted coexistence – all else being equal, communities with higher intraspecific competition tended to retain a higher fraction of species following a perturbation (Fig 2B). Note that this result also indicates that communities with a large fraction of cooperative interactions may still be stable if intraspecific competition is sufficiently high (Barabás et al., 2017; Bascompte and Scheffer, 2023). Following these extinctions, the resultant community is stable, and a second perturbation did not lead to any major changes in the community composition. This is demonstrated by the fact that the number of species in the community following one perturbation was not significantly different from that of species following two perturbations (Fig 2C. Wilcoxon rank-sum text, *W = 5193, p* = 0.63). Broadly the same results hold for communities with exploitation (Figs S1, S2). To assess how the community composition changes after a fraction of the species is lost, we studied whether the proportion of interactions of each type in the post-perturbation community significantly differed from that of the original community.

**Figure 2:**
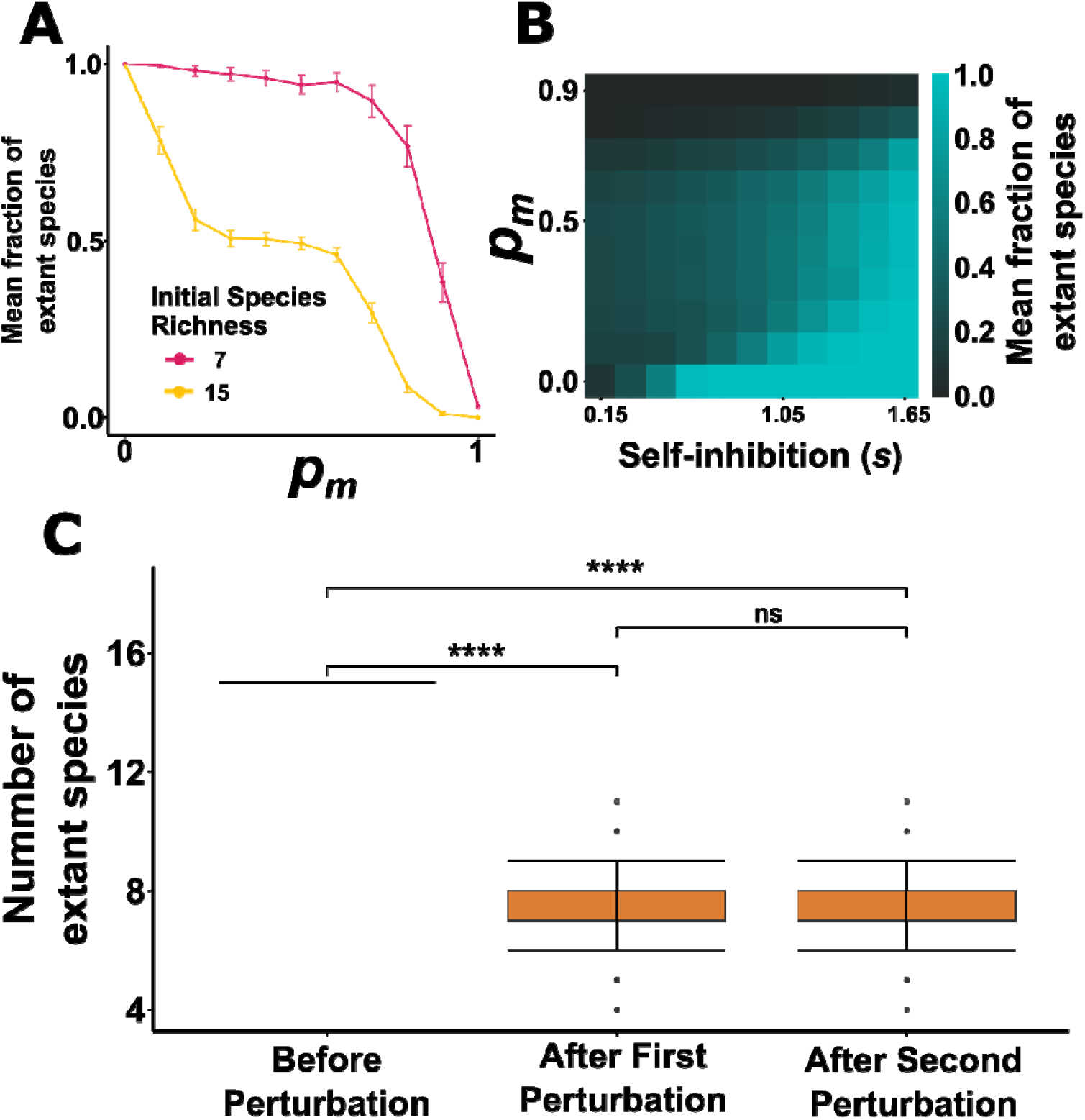
Fraction of extant species left after perturbation. In all simulations, interaction strengths are drawn from a Gamma distribution such that the mean is 0.25 and the variance is 0.01. **(A)** For a given strength of intraspecific competition (here, s=1.05), the mean fraction of extant species reduces with increasing but takes non-zero values for most values of The fraction of extant species is higher for a smaller community (here, seven species) compared to a larger community (here, 15 species). Each point is the mean of 100 realizations, and error bars represent 95% CIs. **(B)** Mean fraction of extant species (averaged over 100 realizations) increases with strength of intraspecific competition across different values of for a community of 15 species. **(C)** The number of extant species present in an unstable community (S = 15, = 0.5, s = 1.05) falls after one perturbation, but does not change following subsequent perturbations. (Wilcoxon rank-sum test, n=100, W = 4892.5, p > 0.1).

### Competitive Interactions are selectively lost following a perturbation in unstable communities

If all species have equal probabilities of going extinct after a perturbation, then, on average, we expect the proportion of competitive (or cooperative) interactions in our randomly assembled communities to remain unchanged by the end of the simulation. Our results reveal that this is not the case. The stable communities formed after species loss had a larger fraction of cooperative interactions and a lower fraction of competitive interactions than the initial starting communities (Fig 3A). In other words, if some species go extinct in a community following a perturbation, the overall amount of competition in the community, measured in terms of the fraction of competitive interactions, tends to reduce. In communities without exploitation, this implies that despite cooperation having been associated with decreased stability in earlier studies, the fraction of surviving cooperative interactions would be higher than in the original community prior to perturbation.

**Figure 3:**
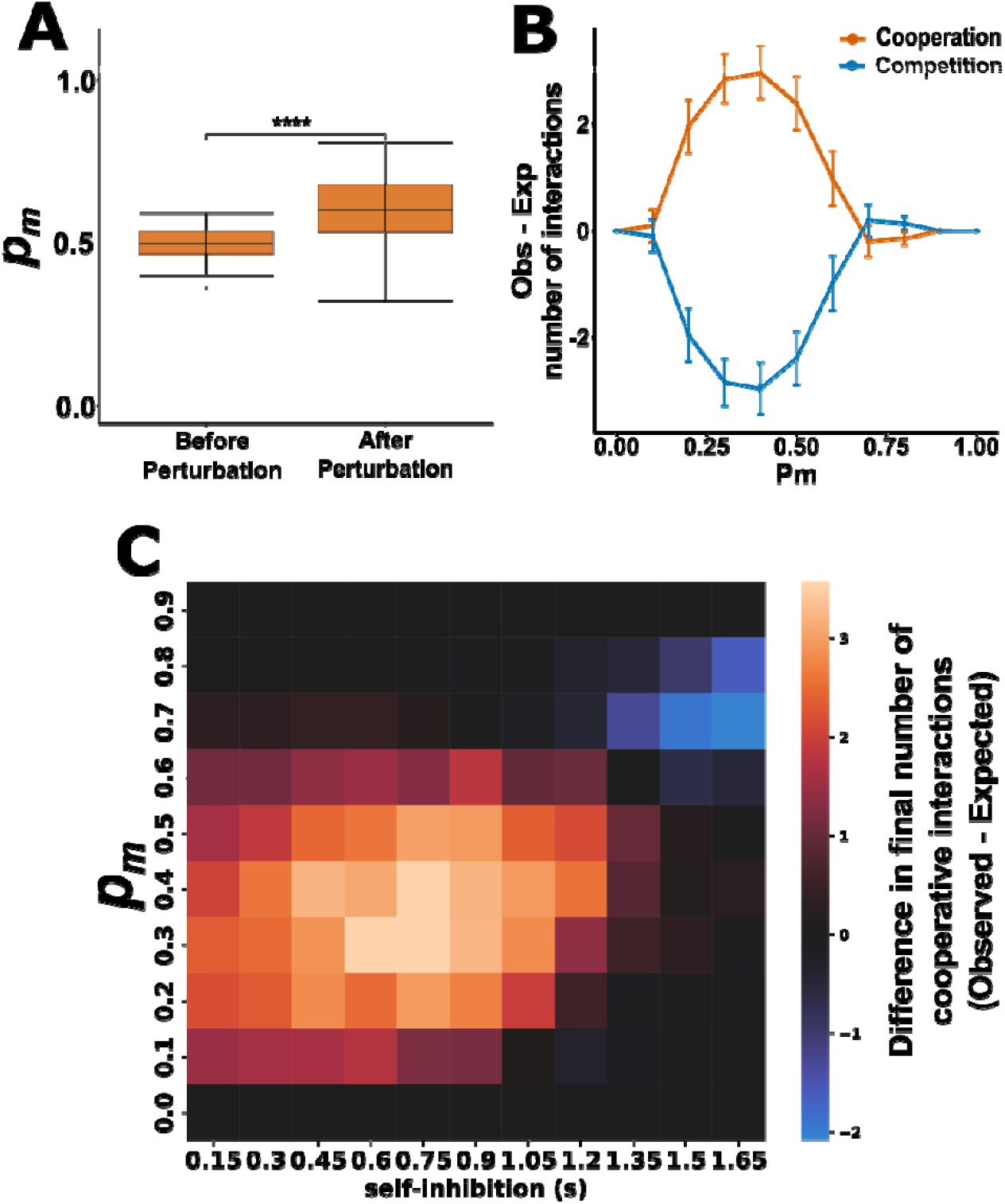
The fate of cooperative and competitive interactions after species loss. In all simulations, S=15 before perturbations. Interaction strengths are drawn from a Gamma distribution such that the mean is 0.25 and the variance is 0.01. By plotting the difference between the observed and expected values of the number of interactions of each type, we can examine whether some interaction types tend to be preserved after extinction events. Here, the expected values are the values of the original community, while the observed values are those measured after equilibrium attainment following the induced perturbation. If this value is greater than 0, fewer interactions of that type are being lost than would be expected by chance alone, and if the value is less than 0, then more interactions are being lost than expected by chance alone. **(A)** The difference in the proportion of interactions before and after a perturbation is statistically significant, as tested by a Wilcoxon rank-sum test (n = 100, W = 1737, p < 0.001, effect size r = 0.564). In this plot, s =1.05. **(B)** For a fixed value of s (here, s = 1.05), regardless of the value of p_m_, on average, cooperative interactions tend to be lost less often than expected by chance alone. In contrast, chance alone loses competitive interactions more often than expected. In this plot, the points represent the mean over 100 realizations, and error bars represent 95% CIs. **(C)** This qualitative result is valid for a large fraction of the parameter space, as indicated by heatmaps in which p_m_ is varied along the y-axis and s is varied along the x-axis. In these plots, the color represents the mean difference (over 100 realizations) between the observed and expected number of interactions following a perturbation. Warmer / Redder colors indicate that the difference is greater than zero (more cooperative interactions are retained), and cooler / bluer colors indicate that the difference is less than zero (i.e., more competitive interactions are retained). For a large region of the parameter space, cooperation tends to be preferentially retained, whereas competition tends to be preferentially lost.

Indeed, our results indicate that communities found after perturbation have a significantly higher fraction of cooperative interactions, as measured by a Wilcoxon rank-sum test (Fig 3A, W = 1737, *p* < 0.001), and effect size calculations indicate that this bias has a large effect (Wilcoxon effect size *r* = 0.564). This broad result is valid for a large array of (Fig 3B, 3C) and intraspecific competition (Fig 3C) values, suggesting that the result is robust to variation in initial community composition. Thus, even though ‘more cooperative communities’ (i.e., communities with a greater fraction of cooperative interactions between species) are less stable, perturbation-driven-extinctions in these communities do not lead to communities with a lower proportion of cooperative interactions. However, at high *p*_*m*_ and high intraspecific competition, the trend is reversed and, albeit for a small fraction of the parameter space, competitive interactions are lost less often than expected by chance alone (Fig 3C). Broadly the same results hold for communities with exploitation, except that it is exploitation that is lost less often than expected by chance alone, whereas both cooperation and competition may be lost either more often or less often than expected by chance alone (Figs S3-S7). Though competitive interactions may be lost either more or less often than expected by chance alone in communities with exploitation, the magnitude of the bias away from chance expectations is greater when competition is selectively lost (Fig S6). The result presented in Fig 3C also holds for communities with smaller species richness (*S=7*), albeit with a smaller magnitude of the effect (Fig S8,S9).

The greater loss of competitive interactions can be understood through the following reasoning. Given any two species and, the net effect of species on the growth rate of species depends on both their interaction type and strength () and on the population density of species (). Thus, for a given species, if the magnitude of interspecific interaction strengths () are approximately similar, the strongest interspecific effect will be from that species with the highest population density (). In other words, if interaction strengths are similar, the most significant interspecific effect on species *i* will be caused by the species *j* which has the largest number of individuals. Immediately following a perturbation (*i*.*e*., random killing of a fraction of the individuals in the community), highly competitive species are likely to experience a larger growth rate, and species that are more cooperative are likely to experience a lower growth rate. For two species engaged in a cooperative (+/+) interaction, the reduction of population size of one species (let’s call it the target species) leads to a reduction in the realized growth rate of the other species (let’s call it the partner species). Since, in our simulations, the communities begin at equilibrium (*i*.*e*. zero growth rate), any reduction makes the growth rate negative, thus reducing the population size of the partner species. This in turn causes a decrease in population size of the target species by the same logic. Thus, cooperative species tend to drive themselves to ever-lower numbers through positive feedback loops (Coyte et al., 2015; Lever et al., 2020).

Species which are engaged in a large number of competitive interactions enjoy a release from competition immediately after a perturbation and thus are likely to have the largest population size in the community. In Fig 4a, we show that the species engaged in the most number of competitive species, which we call the ‘top competitor’, quickly becomes the dominant species in the community following a perturbation (Fig 4a, black trajectory). However, since interactions are assigned at random, even the top competitor will likely be engaged in a (small number of) cooperative interactions. Once a small number of such competitive species have risen to relatively large population sizes, they will then ‘pull up’ those species which are engaged in cooperative interactions with them (Fig 4A, red trajectories) while ‘pushing down’ other competitors and driving them closer to extinction (Fig 4A, grey trajectories). Thus, species engaged in positive interactions with highly competitive species tend to go extinct less often than expected by chance alone (Fig 4B. Wilcoxon Rank-sum test, *W* = 6113.5, *p* < 0.001), and effect size calculations indicate that the reduction in extinction rate is moderately large (Wilcoxon effect size *r = 0*.*338*). Since we study random, unstructured communities, species engaged in a higher number of cooperative interactions are more likely to be engaged in cooperative interactions with the top competitors. In contrast, species engaged in a higher number of competitive interactions are more likely to be engaged in competitive interactions with the top competitors (probabilistically). Thus, species engaged in more cooperative interactions are more likely to get a boost in growth rate due to being engaged in positive interactions with highly competitive species. On the other hand, more competitive species are likely to be engaged in competitive interactions with the top competitor and are thus pushed to extinction. This mechanism does not work if the proportion of cooperation is very high because all competitors, in this case, are likely supported by a large number of cooperative interactions, which explains the trend reversal at high *p*_*m*_ values. This explanation reveals that species engaged in a large number of exploitative interactions enjoy two distinct benefits following a perturbation --- species that are exploited by many species obtain a large increase in growth rate immediately following a perturbation due to release from exploitation/predation, whereas species that exploit a large number of species experience an increase in growth rate through a mechanism similar to that explained for mutualists above. Indeed, in communities with exploitative interactions, simulations reveal that the proportion of exploitative interactions always increases following a perturbation (Figs S4, S7).

**Figure 4:**
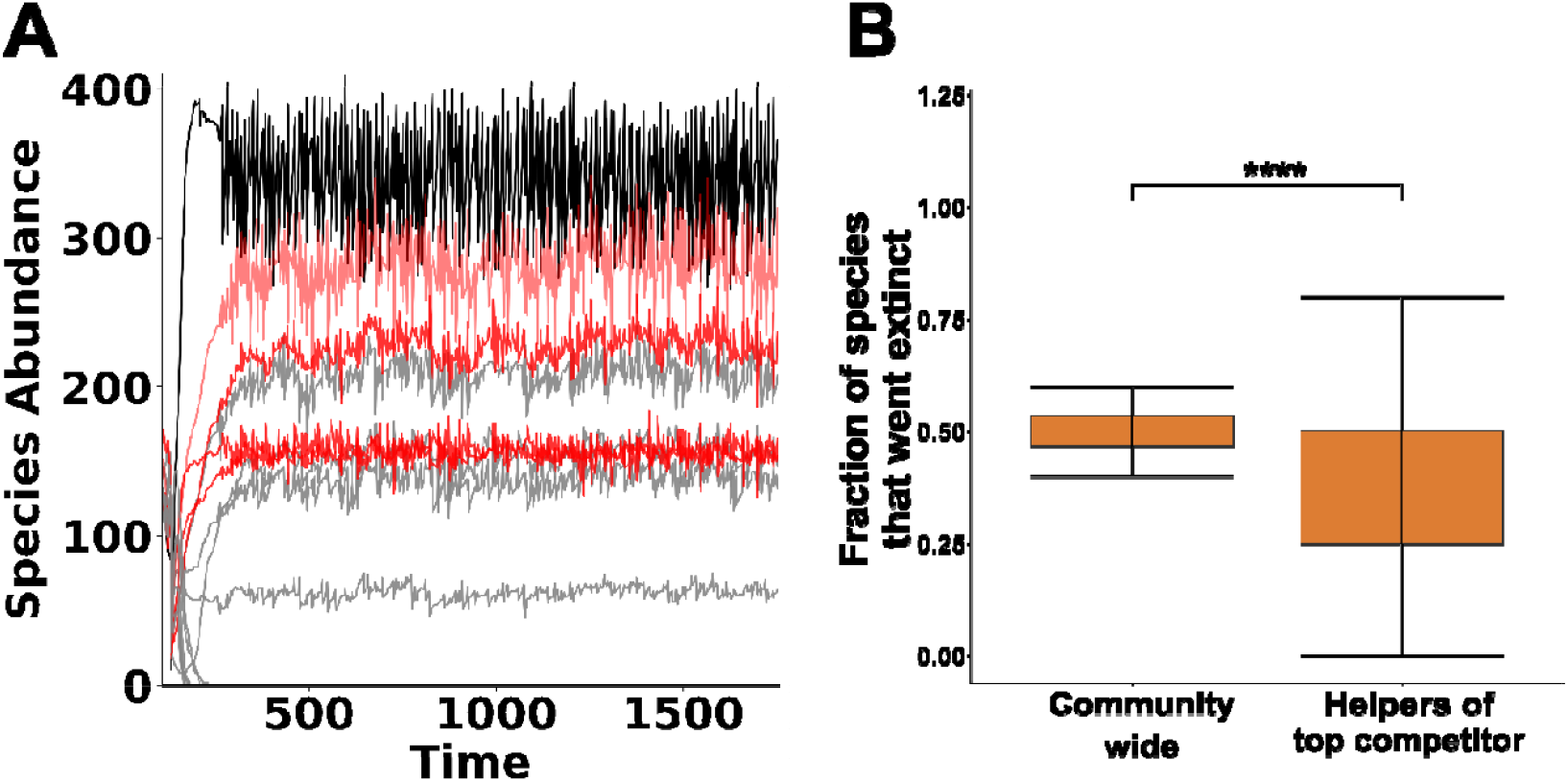
Mechanism for the stability of communities induced by extinctions in unstable communities. **(A)** The trajectories of individual populations of each species for a single realization of the IBM with S = 15, p_m_ = 0.5, s = 1.05 are plotted. The species with the largest number of competitive interactions (the ‘top competitor’) is colored in black, species engaged in cooperative interactions with the top competitor are colored in red, and all other species are colored in grey. Following a perturbation, the top competitor quickly increases in numbers and ‘pulls along’ those species engaged in cooperative interactions while driving the others to extinction. **(B)** Statistical analysis of 100 independent realizations reveals that for this set of parameters (S = 15, pm = 0.5, s = 1.05) if the top competitor does not go extinct, the extinction probability of species that are engaged in cooperative interactions with the top competitor is significantly less than the background extinction probability of the community as a whole. (Wilcoxon rank-sum test, W = 6945.5, p < 0.0001, effect size r = 0.338). The center of the box plot denotes the median, and the edges of the box indicate the upper and lower quartile.

Our explanation of how competitive interactions are selectively removed while cooperative interactions are maintained hinges on the assumption that the community is unstructured and can be modeled as a random matrix of interactions between species. Communities in nature, of course, may not necessarily satisfy these criteria. For example, there may be a community where the most competitive species do not engage in any cooperative interactions. In such a community, even though some highly competitive species may see a sharp rise in population density following a perturbation, they may not pull up the numbers of species that participate in cooperative interactions. Our analysis does not attempt to model such exceptional cases and is meant to interpret the results for typical random, unstructured communities. Strictly speaking, exceptional cases, such as those mentioned above, can arise despite a random assignment of interactions between species. However, the probability of such communities in our design is extremely low and therefore are unlikely to play a critical role in driving the general trends.

## Discussion

Our results indicate that when a community is unstable (in the persistence sense, i.e., likely to lose species after a perturbation), only a subset of the species in the community goes extinct before the community becomes stable again (Fig 2C), recapitulating the results of Rohr et al 2025 using an individual-based model. Furthermore, we show that species engaged in more cooperative interactions are less likely to go extinct, suggesting that the extinction patterns in randomly assembled communities are non-random with respect to interaction type. While similar results predicting that competitive interactions are more likely to go extinct have been presented for food webs (Barabás et al., 2017) and structured communities with nested interactions (Thébault and Fontaine, 2010), our results indicate that unstructured (*i*.*e*. randomly assembled) communities with multiple types of interactions also exhibit the same bias towards losing competitive interactions in the event of an extinction. The effects we uncover thus represent systematic biases in extinction probability that will consistently affect resultant community dynamics following any external perturbation that is strong enough to lead to species loss. Together with previous results (Thébault and Fontaine, 2010, Barabás et al., 2017), these dynamics suggest a potential explanation for the prevalence of cooperation in natural communities (Kehe et al., 2021; Machado et al., 2021) – a community can harbor reasonably high levels of cooperation if it is formed due to species loss from a larger randomly interacting community. Community assembly is often thought to occur by random dispersal followed by environmental filtering and subsequent exclusion of some species due to biotic interactions (Begon et al., 2006; Molles, 2015). These are precisely the kind of processes for which our results would be relevant. Thus, our model highlights the importance of assembly processes in determining community structure.

A recent modeling study of assembly processes in communities also suggests that when species sequentially invade a community, a balance of interaction types is vital for community stability, with higher fractions of cooperative interactions corresponding to increased species persistence as well as increased stability of the community as a whole to external invasions (Qian and Akçay, 2020). Such so-called ‘ecological selection’ (Qian and Akçay, 2020) for community structure during assembly has also been observed in dispersal models (Denk and Hallatschek, 2023) and eco-evolutionary community models ((Serván and Allesina, 2021); Nell et al., 2022). Our study highlights that ecological selection of this form can operate not only through a sequential assembly of communities but also through extinctions from initially assembled unstable communities. In nature, a situation mirroring our model is often encountered in communities such as microbiomes, where empirical data suggests that many species are randomly assembled through dispersal processes and environmental filtering (Sieber et al., 2019; Venkataraman et al., 2015). Empirical studies of microbial communities also suggest that positive interactions, in the sense of exploitations as well as mutualisms, are ubiquitous in culturable bacteria (Kehe et al., 2021) (but see (Palmer and Foster, 2022)). Our study provides a mechanistic hypothesis for reconciliation of such empirical results with theory regarding the destabilizing effects of cooperation (Allesina and Tang, 2012; Coyte et al., 2015).

Our model also predicts an emergent non-random interaction structure from an initially unstructured, unstable community. Since competitive interactions with the top competitor are selectively lost, the stable community so formed is likely to have a small number of ‘central’ species (previously the ‘top competitors’) engaged in positive interactions with most community members. Competitive interactions with this ‘central’ species should be relatively weak. This aligns with a previous analytical study, which predicts that assembly processes should lead to ecological networks with weaker competition and stronger cooperation than the original species pool (Bunin, 2016). A similar phenomenon has also been observed in an analytical model of unstructured communities with higher-order interactions (Gibbs et al., 2022). In this study, the authors found that when interaction strengths in their model were low, the dominant species in communities with higher-order interactions tended to be those engaged in positive interactions with each other and engaged in negative interactions with species that have lower species density. By studying the dynamics of unstable communities, our results underscore the need to go beyond the question of whether communities are stable to study the fate of unstable communities. While such studies are often complex to conduct analytically, computational methods, laboratory experiments, and long-term field observations provide potential avenues to address this vital question.

Though we have only looked at randomly assembled communities, a previous simulation study (García-Callejas et al., 2018) suggests similar results may hold for structured trophic networks. These authors found that positive interactions such as cooperation and commensalism promoted persistence in trophic networks with low species richness. However, this effect was less pronounced at higher species richness. Our study neglects environmental or spatial heterogeneity, which is known to affect coexistence and stability (Allen et al., 2013; Durrett and Levin, 1994; Gordon et al., 2015; Hauert and Doebeli, 2021; Krakauer and Pagel, 1995; Stein et al., 2014; Ursell, 2021; Yu et al., 2001). Another factor that can potentially affect community stability is demographic stochasticity, which has been shown to promote cooperation in many model systems (Chotibut and Nelson, 2015; Constable et al., 2016; Houchmandzadeh, 2015; Houchmandzadeh and Vallade, 2012; McLeod and Day, 2019; Altieri et al., 2021). Lastly, non-linearities in population dynamics can also stabilize community population dynamics and favour coexistence, especially in mutualistic networks (Bascompte and Scheffer, 2023; Bastolla et al., 2009; Holland et al., 2002; Lever et al., 2020; Ringel et al., 1996). By neglecting these factors, we do not mean to imply that they are unimportant. Instead, we illustrate that they cannot *solely* be responsible for explaining the occurrence of cooperation since, as we have shown, cooperation may persist purely through non-random extinction processes during initial community assembly.

We also do not study the possibility of targeted perturbations that only affect certain particular species in a community. The positive feedback loop proposed by Coyte et al. 2015 for the destabilizing effect of cooperative species suggests that selectively reducing the population of the most cooperative species in the population should lead to other cooperative species in the community going extinct. The mechanism we propose for the selective loss of competitive interactions following a perturbation (Fig 3C and Fig 4) also suggests that selectively perturbing the density of the top competitor should lead to an increase in the densities of all other species engaged in competitive interactions with this top species. However, a complete investigation into the effects of non-random perturbations is beyond the scope of the present study and may prove an interesting direction for future work.

Our model also assumes that all direct community interactions are pairwise and allows higher-order interactions to only manifest as emergent properties of the simulation. Analytical studies suggest that many classic results from pairwise interaction models carry over to models with higher-order interactions (Gibbs et al., 2022). In particular, May’s classic results on the diversity-stability relation carry over to models with higher-order interactions (Gibbs et al., 2022). Therefore, there is a possibility that such models may lead to results analogous to those presented in this study. However, it bears noting that a large number of species can be made to coexist in a community if higher-order interactions are tuned in a non-random way to enhance effective competition strengths. In particular, if higher-order interactions in a community are structured such that the interactions overall strongly increase the effective intraspecific competition (relative to effective pairwise interspecific competition), May’s result can be inverted and species richness can promote stability (Grilli et al., 2017; Singh and Baruah, 2021). Structured interactions can also favour mutualism more generally if the community structure is wired to be such that the effective intraspecific competition is high (Bascompte and Scheffer, 2023; Bastolla et al., 2009; Lever et al., 2020; Ringel et al., 1996).

A recent paper has argued on analytic grounds that cooperation also does not reduce resilience in Lotka-Volterra communities if studies use the so-called ‘community matrix’, which differs from the more commonly used ‘interaction matrix’ by accounting for species densities (Stone, 2020). Our explanation for the non-random loss of competitive interactions from an unstable community following a perturbation also crucially relies on the observation that species densities play a large role in determining community dynamics. However, in contrast to the analytical study, our model predicts a decrease in stability with an increase in cooperation, but nevertheless, provides a potential explanation for the prevalence of cooperative interactions in nature, namely ‘ecological selection’.

Lastly, our simulations are over ecological timescales and do not allow for evolution via speciation or evolution in traits (interaction strengths and/or intrinsic growth rates). Incorporating evolution is known to qualitatively alter the predictions of purely ecological models (Kokko et al., 2017; Schoener, 2011; Yamamichi et al., 2022). A suite of empirical studies increasingly indicates that the separation of timescales between ecology and evolution can often be blurred, especially in the case of organisms such as microbes (Schoener, 2011). Including evolutionary processes in ecological coexistence theory and the cooperation-competition debate thus provides an attractive avenue for future studies.

## Acknowledgments

The authors would like to acknowledge Akshay Malwade for several helpful discussions. Two anonymous reviewers provided helpful comments that greatly improved an earlier draft of this manuscript. The support and the resources provided by the PARAM Brahma Facility under the National Supercomputing Mission, Government of India, at the Indian Institute of Science Education and Research (IISER) Pune, are gratefully acknowledged. ASB and SN are supported by Kishore Vaiyanik Protsahan Yojana (KVPY) fellowships from the Department of Science and Technology, Government of India. This project was supported by grant # STR/2021/000021 from Science and Engineering Research Board, Department of Science and Technology, Government of India and internal funding from Indian Institute of Science Education and Research, Pune.

## Author contributions

**ASB**: Conceptualization, Methodology, Software, Formal analysis, Investigation, Writing – Original Draft, Writing – Review & Editing, Visualization; **SN**: Conceptualization, Methodology, Formal analysis, Investigation, Validation, Writing – Original Draft, Writing – Review & Editing; **SD**: Conceptualization, Methodology, Resources, Writing – Review & Editing, Supervision, Funding Acquisition.

## Data archiving statement

All codes and data used in this manuscript will be available online after acceptance.

