## Supplement for "Cooperation Destabilizes Communities, but Competition Pays the Price"

**Supplementary Information**

This document contains supplementary figures for the study ‘Mutualism destabilizes communities, but competition pays the price’. The main text contains the results of simulations in which *p_e_ = 0* (*i.e.* all interspecific interactions are either cooperative or competitive). In figures S1-S7, we present the corresponding plots for communities in which exploitation is present in equal proportion as competition (*i.e. p_c_ = p_e_*). As in the main text, all means are over 100 independent realizations, and in plots with varying *p_m_, p_m_* is varied from 0 to 1.

In Figures S8 and S9, we show that the result presented as Fig 3C in the main text also holds for communities with a smaller species richness (*S = 7*), albeit with a smaller magnitude of the effect.

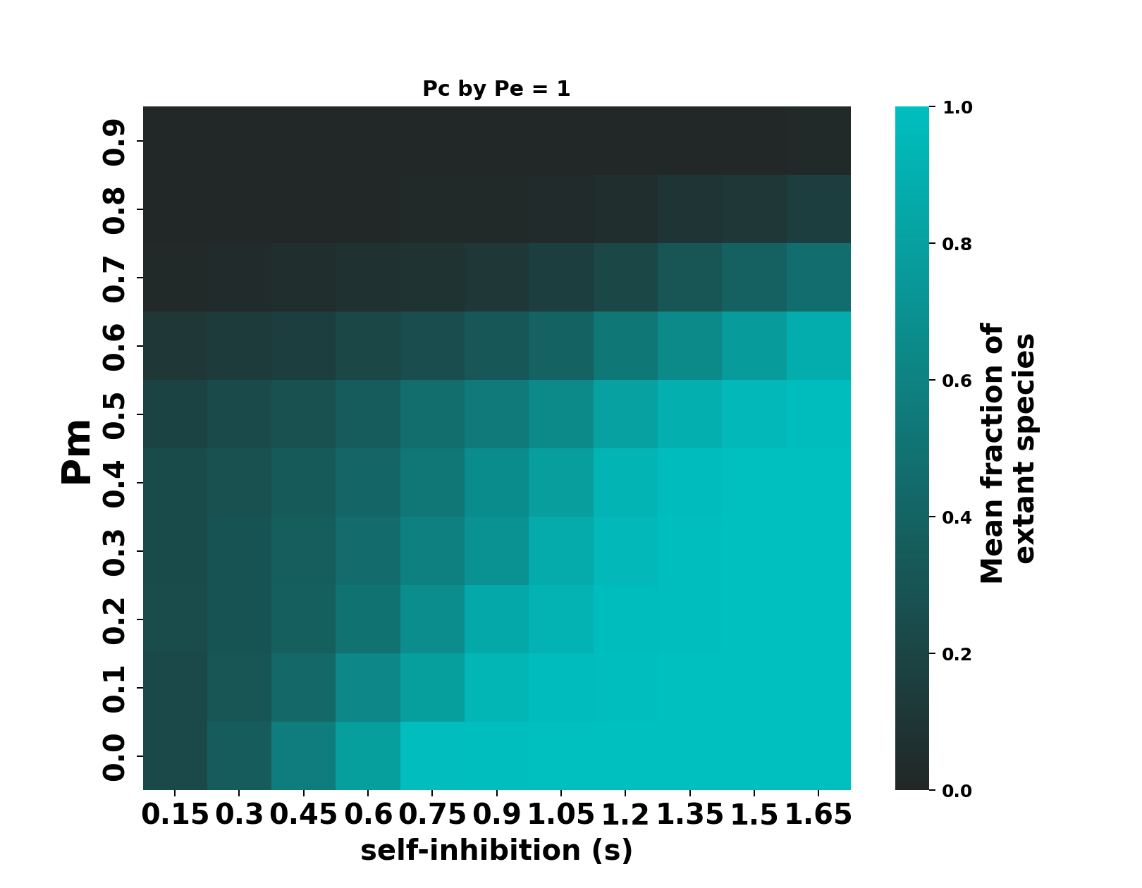

*Fig S1: Proportion of extant species as a function of amount of mutualism (p­_m_) and intraspecific competition (s). Compare with Fig 2B in the main text.*

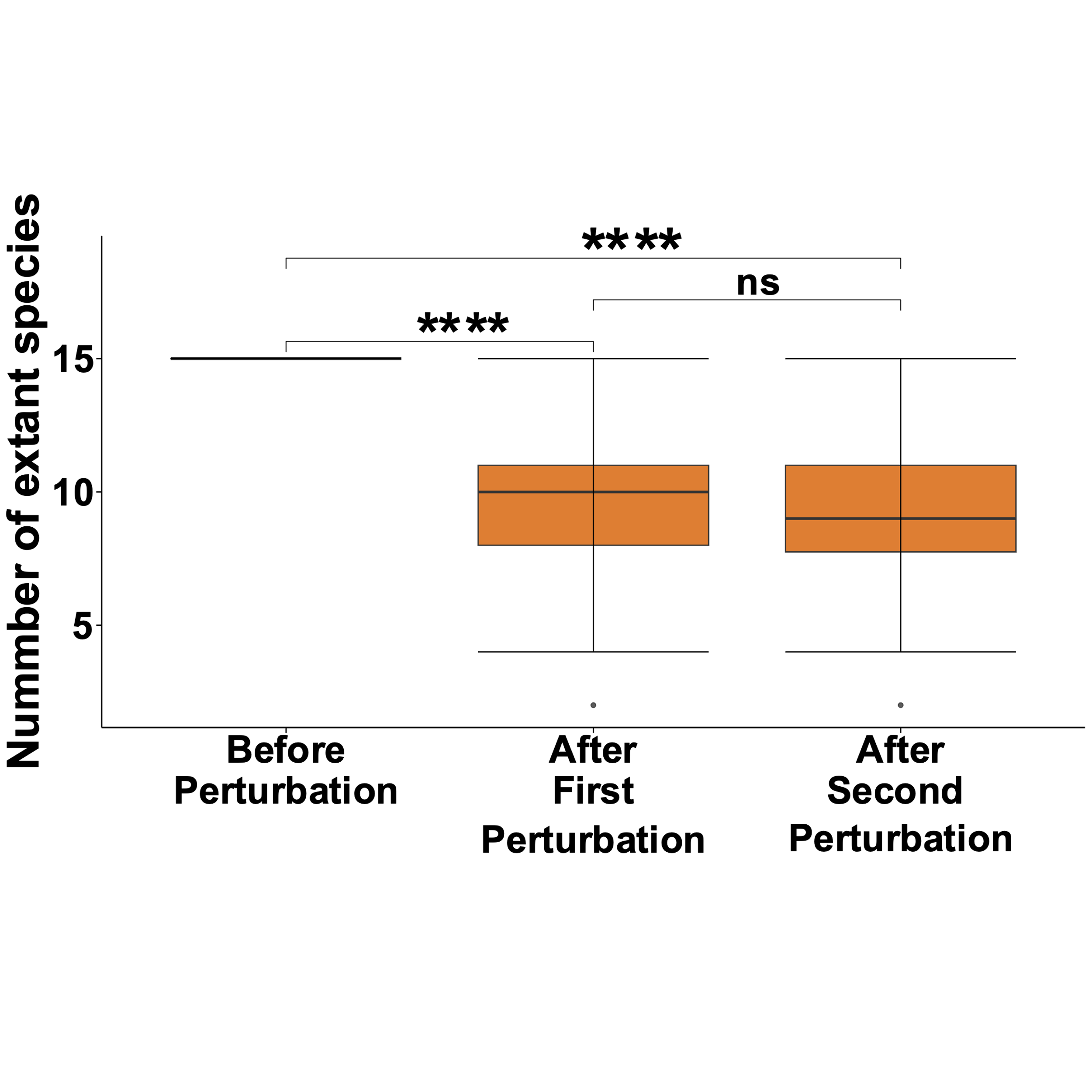

*Fig S2: Proportion of extant species following one perturbation and two perturbations for communities in which p_c_ = p_e_. Following a perturbation, communities experienced significant species loss (Wilcoxon rank sum test, W=9700, p < 0.0001). However, the number of species left extant in the community following a single perturbation was not significantly different from that after two perturbations (Wilcoxon rank sum test, W = 5559.5, p = 0.17). Compare with Fig 2C in the main text. Legend is same as for Fig 2C. Here, s = 1.05 and p_m_ = 0.5.*

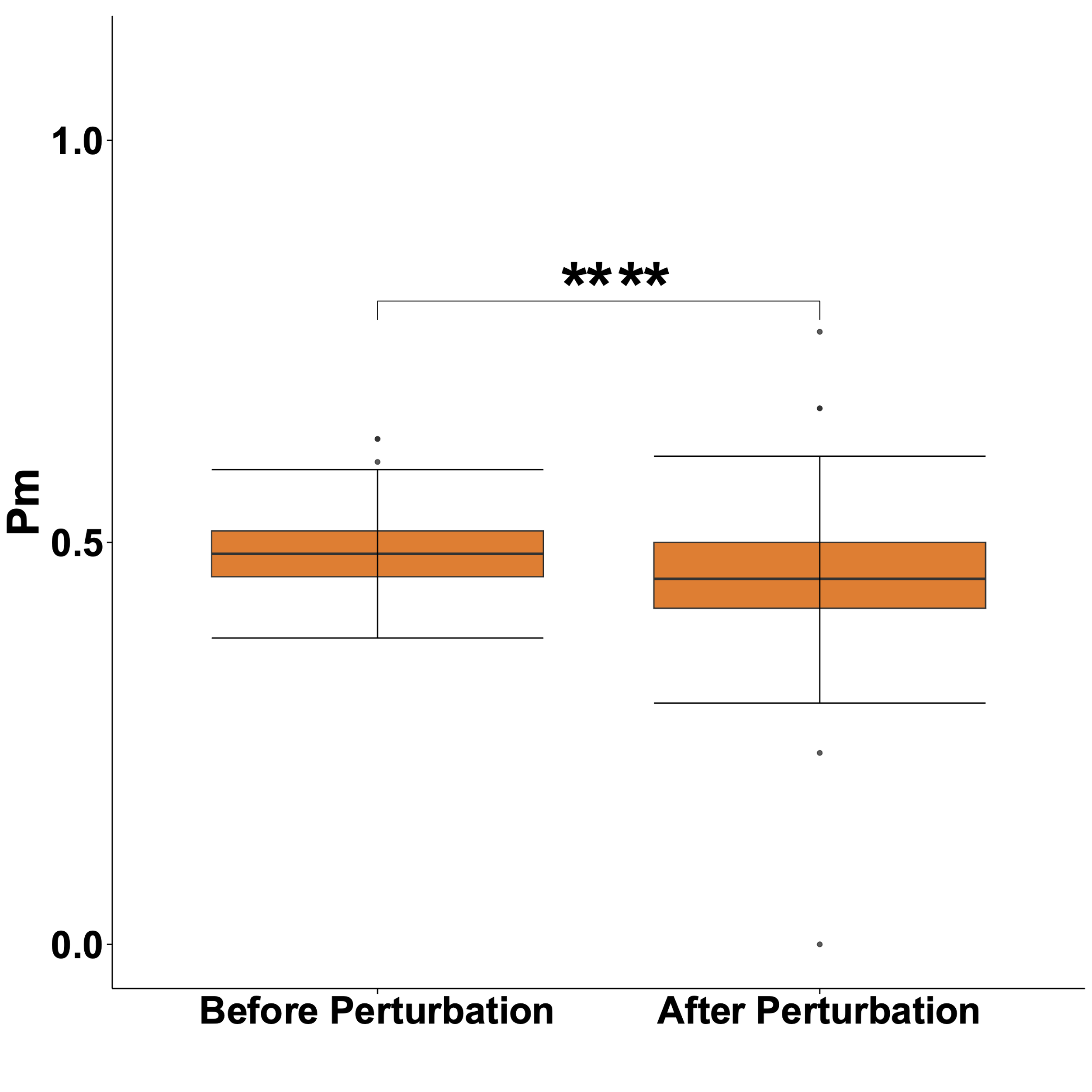

*Fig S3: Proportion of cooperative interactions before and after perturbation for communities in which p_c_ = p_e_. In this case, p_m_ slightly decreases (Wilcoxon rank sum test, W = 6594.5, p < 0.0001). However, the effect is small (Wilcoxon effect size r = 0.276) and is not a general trend, as will be revealed upon examining the full parameter space in Figs S5-S7. Here, s = 1.05. Compare with Fig 3A in the main text. Legend is same as for Fig 3A.*

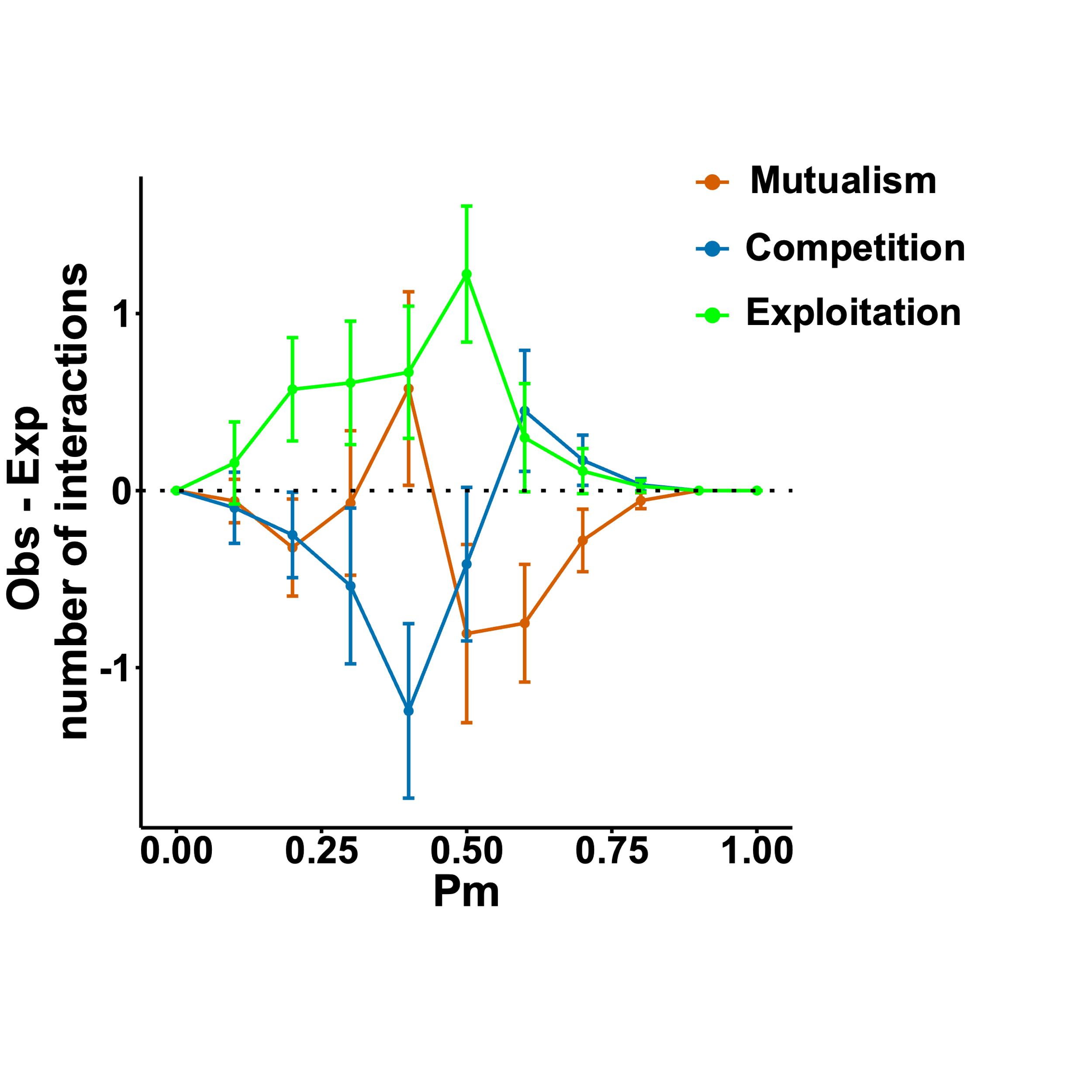

*Fig S4: The difference between observed and expected number of interactions of each type for communities in which p_c_ = p_e_. In this case, the dynamics are more complicated than those of communities with no exploitation. Broadly, exploitation is always lost less than expected by chance alone, whereas both mutualism and competition may be lost either more often or less often than expected by chance alone, though in opposite regions of parameter space. In this plot, the intraspecific competition strength is s = 1.05. Error bars represent standard errors, and points are mean values. Compare with Fig 3B in the main text.*

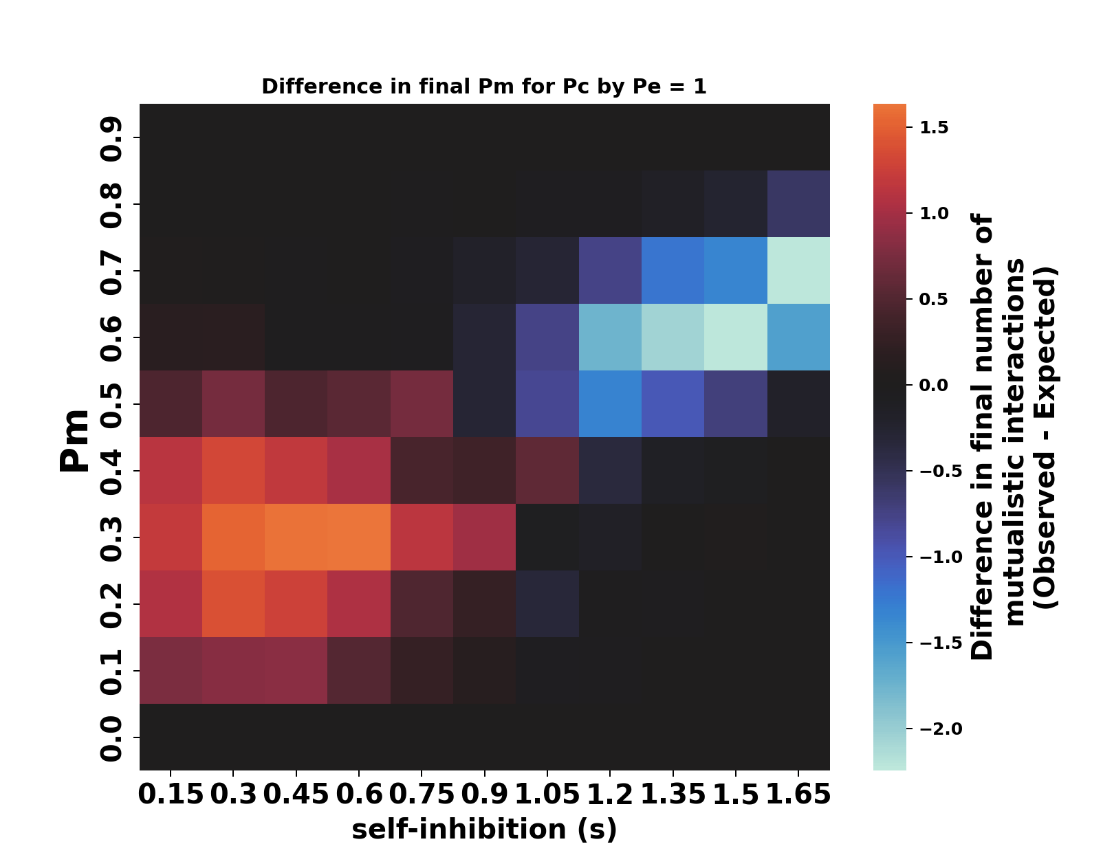

*Fig S5: Observed vs expected number of cooperative interactions following a perturbation. In communities with exploitation, there is also a significant range of parameter space in which mutualism is lost more often than expected by chance alone. However, the effects are generally rather weak (compare limits of color bar with those of Fig 3C in the main text). Legend is same as in Fig 3C.*

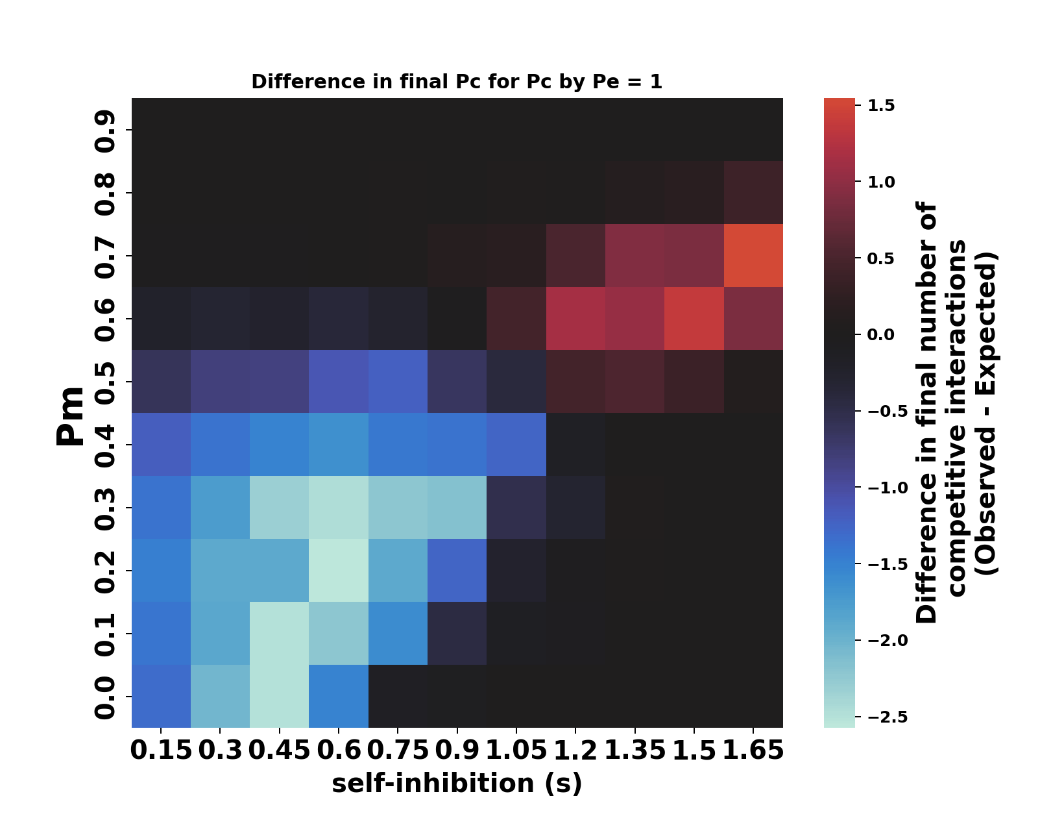

*Fig S6: Observed vs expected number of competitive interactions following a perturbation. Competition may be lost less often or more often than expected by chance alone. However, the gain in competition is weaker in effect than the loss (compare red limit vs blue limit in color bar). Legend is same as in Fig 3C.*

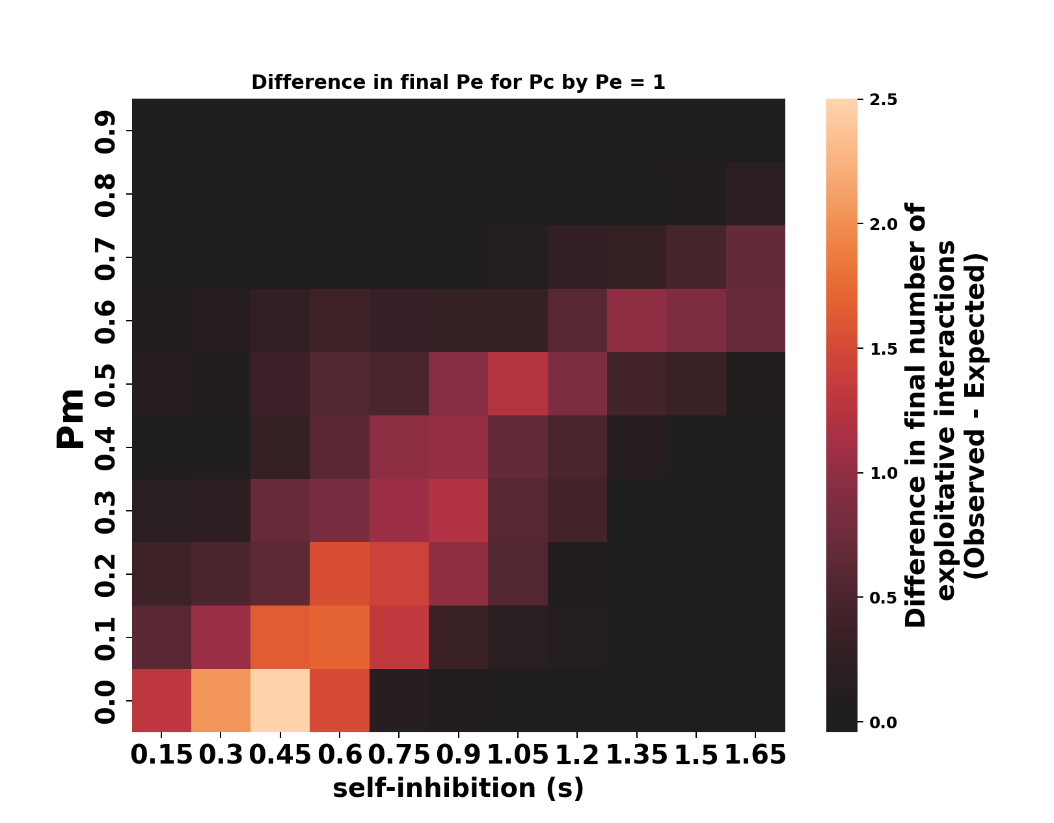

*Fig S7: Observed vs expected number of exploitative interactions following a perturbation. Exploitative interactions are always lost less often than expected by chance alone. Legend is same as in Fig 3C.*

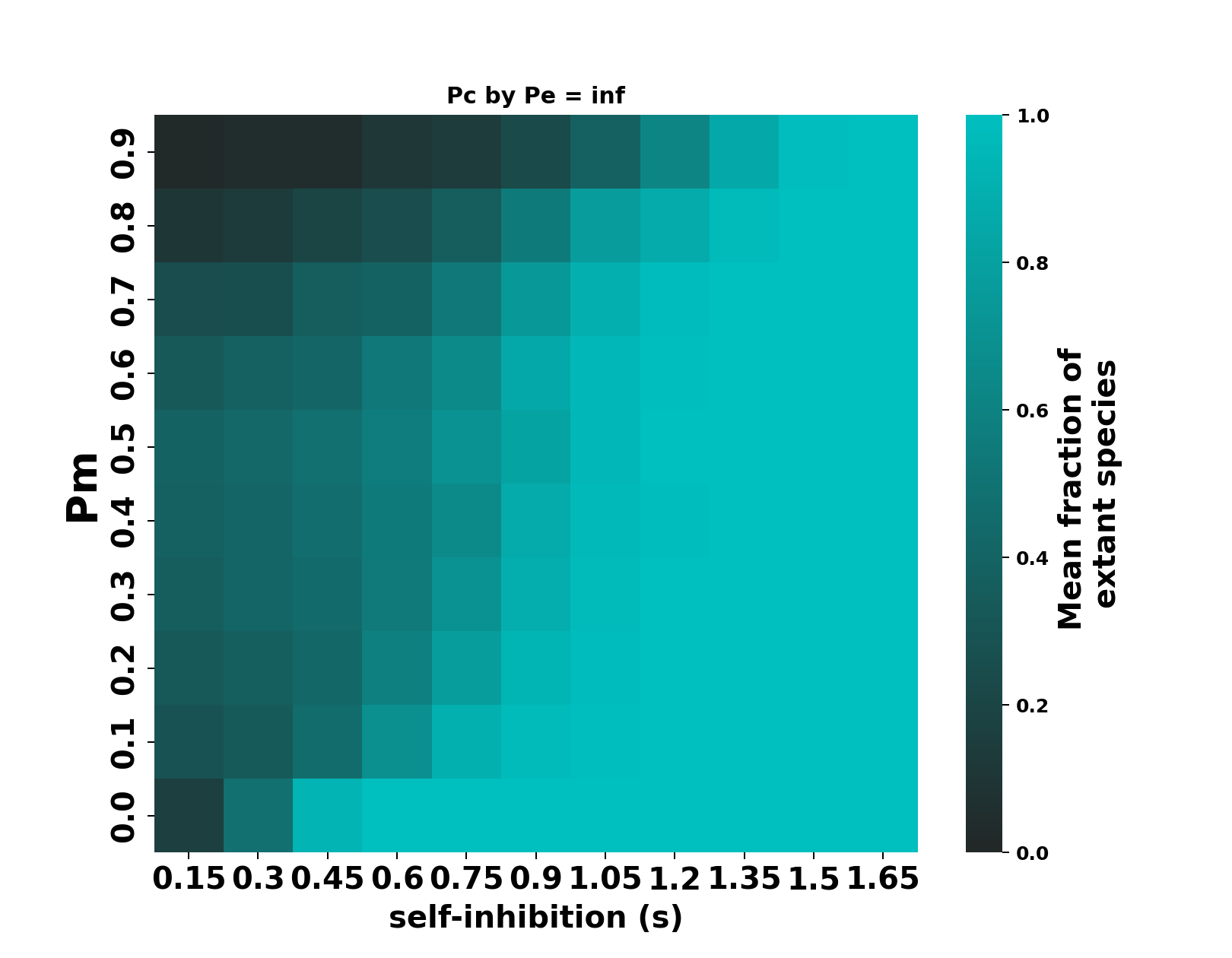

*Fig S8: Fraction of extant species following a perturbation as a function of p_m_ and self-inhibition in a community consisting of 7 species (S=7). Compare with Figure 2B in the main text. Legend is same as in Fig 2B.*

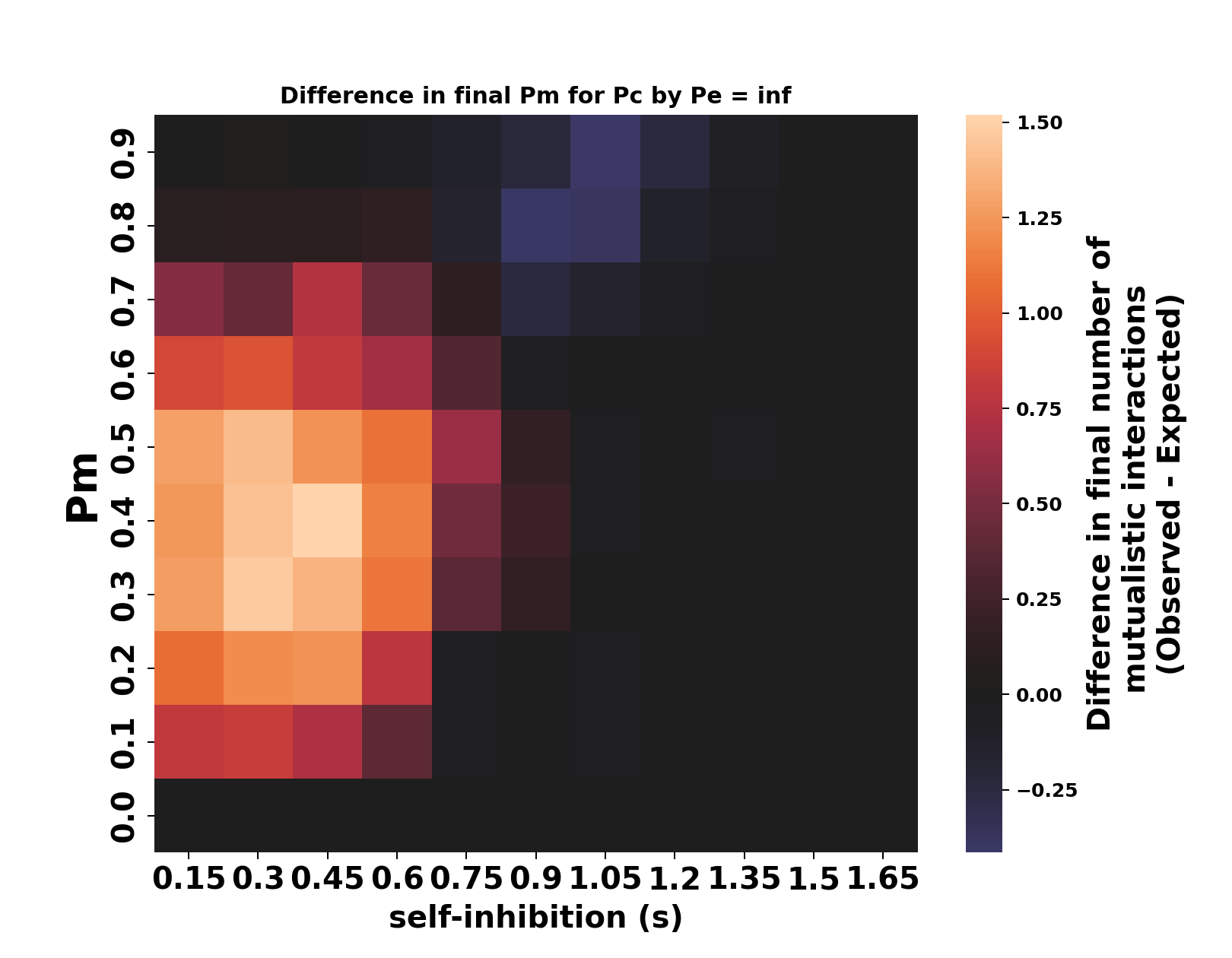

*Fig S9: Observed vs expected number of cooperative interactions following a perturbation in a community with 7 species* (*S = 7*) *in which every interaction is either cooperative or competitive (i.e. p_e_ = 0). Cooperative interactions are thus lost less often than expected by chance alone, whereas competitive ones are lost more often than expected by chance alone, just as in the main text. Legend is same as in Fig 3C.*
